# Conserved cold tolerance of *Rhagoletis* species from different host fruits, elevations in Colorado, USA

**DOI:** 10.1101/2023.12.22.573084

**Authors:** Katelyn Lemay, Mackenzie Moore, Paige Brown, Lahari Gadey, Gregory J. Ragland, Jantina Toxopeus

**Author notes:** Corresponding author; JT. **Emails**: KL -; MM -; PB -; LG -; GJR.

## Abstract

Understanding and characterizing how insects tolerate low temperatures is important for predicting their overwintering survival and subsequent geographic spread. This study characterized the cold tolerance of two members of the *Rhagoletis* genus in Colorado, U.S.A. Pupae were collected from infested fruit in late summer and early fall. For the first time, we show that the rosehip fly *Rhagoletis basiola* is freeze-avoidant; overwintering pupae could supercool to temperatures as low as −26°C and survive. Interestingly, the temperature at which ice forms (supercooling point; SCP) did not vary between *R. basiola* at high (c. 2900 m above sea level) and lower (c. 1650 m a.s.l.) elevations. We also report the apple maggot *Rhagoletis pomonella* infesting an unusual host fruit, the Dolgo crabapple, in close proximity to infested hawthorn trees. *R. pomonella* infesting hawthorn fruits and crabapples had similar SCPs, and survived temperatures as low as −21°C. Pupae from both host fruits also survived prolonged exposure (2 weeks or more) to mild low temperatures (0 to −5°C). Further study into the mechanisms underlying the impressive and conserved cold tolerance of *R. pomonella* and *R. basiola* is an interesting avenue for future research.

## Introduction

Low temperatures associated with winter can pose a significant risk to organisms, especially ectothermic insects. Extreme cold can be lethal for insects, as subzero temperatures can lead to freezing of internal body fluids, potentially damaging tissues and cellular structures (Sinclair et al., 2015; Toxopeus and Sinclair, 2018). Mild cold (e.g., temperatures above or just below 0°C) can also impair organismal function, leading to chill coma (cold-induced paralysis) and chilling-related injury that can be lethal (Overgaard and MacMillan, 2017). The effects of cold depend both on the temperature itself (mild vs. extreme) and the duration of the cold exposure (acute vs. chronic) (Sinclair et al., 2015). Non-lethal cold exposures can be beneficial, in some cases increasing tolerance to subsequent stressors (Teets and Denlinger, 2013); minimizing use of stored energy reserves before, during, or after winter (Hahn and Denlinger, 2011; Roberts et al., 2021; Williams et al., 2015); and facilitating the completion of developmental programs such as diapause (Denlinger, 2022; Feder et al., 1997; Hodek, 1996; Koštál, 2006). In this study, we test the impact of low temperatures on survival and development of two insect species in Colorado – the apple maggot fly *Rhagoletis pomonella* (Walsh) and the rosehip fly *Rhagoletis basiola* (Osten Sacken) – including populations with an unusual host fruit (Dolgo crabapples) and at high elevations (e.g., 2900 m above sea level), respectively.

Understanding and characterizing insect acute and chronic cold tolerance is important for predicting overwintering survival and how insect pests may subsequently spread (Roberts et al., 2021; Sinclair et al., 2015). Cold-tolerant insects have evolved several strategies to mitigate the damaging effects of acute low temperature exposure (Lee, 2010; Sinclair et al., 2015). Freeze-avoidant species can survive extreme low temperatures above their supercooling point (SCP), the temperature at which ice formation begins (Lee, 2010; Sinclair et al., 2015). Conversely, freeze-tolerant species survive freezing (internal ice formation) and temperatures below the SCP (Sinclair, 1999; Sinclair et al., 2015). Chill-susceptible species are not cold-tolerant and are often killed by short exposures to subzero temperatures, well above the temperature at which their body fluids freeze (Overgaard and MacMillan, 2017; Sinclair et al., 2015). Chronic cold tolerance refers to the ability to survive longer cold exposures, an important adaptation for tolerating long winters (McIntyre et al., 2023; Toxopeus et al., 2019). This is usually measured by determining the time that insects can survive at a specific low temperature.

Cold tolerance can vary across species and change based on the time of the year, geographical location, and diet. Many insects in temperate regions increase their cold tolerance in response to autumn conditions, prior to the onset of winter (Sinclair et al., 2015; Toxopeus et al., 2019). For example, many insects enter diapause – a state of developmentally programmed dormancy associated with enhanced stress tolerance – in the fall season (Hand et al., 2016; Koštál, 2006; Wilsterman et al., 2021). Populations that inhabit colder climates (e.g., high latitudes, high elevations) are generally more tolerant of low temperatures than populations and species from warmer climates (Addo-Bediako et al., 2000; Sunday et al., 2011). Diet can also impact cold tolerance by influencing the accumulation of cryoprotective molecules (Koštál et al., 2016, 2011; Li et al., 2014) and storage of energy reserves that are important for survival of prolonged cold exposure (Hahn and Denlinger, 2011). The *Rhagoletis* genus (Diptera: Tephritidae) of frugivorous flies has a wide geographic distribution and feeds on a variety of host fruits, providing a good model system to study potential interactive effects of thermal environment and diet on cold tolerance.

*Rhagoletis* fruit flies infest various plants in Rosaceae and other plant families throughout North America, including economically important fruits such as apples, blueberries, and cherries (Yee et al., 2013, 2015; Yee, 2008). Most *Rhagoletis* spp. are univoltine: adults lay their eggs into host fruit in late summer or early fall, larvae consume part of the fruit, and when the fruit drops to the ground the larvae enter the soil, pupate, and overwinter in diapause for most of the year (Feder et al., 1993; Yee et al., 2013). Some species can lay their eggs in multiple hosts, such as *R. pomonella,* who lay eggs in hawthorn fruits (*Crataegus* spp.) and apples (*Malus domestica*). Some fruits can be infested by multiple *Rhagoletis* species; for example, *R. pomonella, Rhagoletis indifferens, Rhagoletis cingulata* and *Rhagoletis fausta* can all infest cherries (*Prunus cerasus*; (Yee et al., 2013). Studying the impact of cold on both pest and non-pest *Rhagoletis* species is important for understanding their geographic distribution and overwintering capacity.

*Rhagoletis basiola* infests rose plants (*Rosa* spp.) and has a geographical range that expands into colder regions than most other *Rhagoletis* species, but its cold tolerance has not been characterized. For example, *R. basiola* are in the Nearctic region, ranging from southern USA (California and Mississippi) to northern Canada and Alaska (Carroll et al., 2002), and are found at very high elevations (c. 2900 m above sea level in Colorado, USA; see Methods). These fruit flies infest rosehips at a low rate (e.g., < 10% of fruits sampled; (Berlocher and Dixon, 2004; Yee et al., 2015), and are not typically considered pests. Given the distribution of *R. basiola* in locations with extreme winters, we expect *R. basiola* to have high cold tolerance, particularly populations at higher elevations and latitudes. While the geographical distribution (Berlocher and Dixon, 2004; Yee, 2008; Yee et al., 2015) and interactions with host fruits and parasitoid wasps (Averill and Prokopy, 1981; Hoffmeister et al., 2000, 1999; Roitberg and Lalonde, 1991) of *R. basiola* have been partially studied, we know little about its cold tolerance.

*Rhagoletis pomonella* is a well-studied and significant pest in the commercial apple industry (Boller and Prokopy, 1976; Bush et al., 1989; Doellman et al., 2020) that predominantly infests apples (*Malus domestica*) and hawthorn fruits (*Crataegus* spp.). The apple maggot is distributed across western and eastern Canada (except for Newfoundland), throughout 38 of 50 states in the USA, and in the northern and central regions of Mexico (Canadian Food Inspection Agency, 2012; Hood et al., 2013; Rull et al., 2009; Yee and Goughnour, 2006; Yee et al., 2013). Previous work on *R. pomonella* from Illinois and Michigan, USA has established that pupae are freeze-avoidant and survive acute exposures to temperatures above the SCP (c. −20°C; (McIntyre et al., 2023; Toxopeus et al., 2021). *Rhagoletis pomonella* pupae also generally survive prolonged (e.g., 4 months) exposures to mild chilling at 4 - 5°C, as long as they are in diapause (Dambroski and Feder, 2007; Feder et al., 1997; Toxopeus et al., 2021; Yee et al., 2021). In addition, there is an impact of diet on energetics: *R. pomonella* that feed on apples accumulate greater lipid reserves than those feeding on hawthorn fruits (Ragland et al., 2012), which has the potential to influence cold tolerance. However, the impact of host fruit (diet) on cold tolerance has not been examined in *R. pomonella*.

*Rhagoletis* flies that infest different host fruits often exhibit marked, evolved differences in life history timing that could interact with cold tolerance. Apple trees bear fruit earlier in the year than hawthorns, and apple-infesting *R. pomonella* generally eclose as adults, lay eggs and enter diapause earlier in the season than hawthorn-infesting *R. pomonella* (Feder et al., 1997, 1993). This relationship is maintained even when apple and hawthorn populations are overwintered under common conditions in the laboratory (Feder et al., 1993; Smith, 1988) and is driving speciation between sympatric populations of apple- and hawthorn-infesting *R. pomonella* (Dowle et al., 2020; Filchak et al., 2000; Ragland et al., 2017). The duration of cold exposure also impacts developmental timing, with longer chilling of pupae resulting in faster completion of post-chill eclosion, with apple flies eclosing faster than hawthorn flies (Dambroski and Feder, 2007; Feder et al., 1997; Toxopeus et al., 2023; Yee et al., 2023). We have recently observed *Rhagoletis* flies infesting introduced (non-native) crabapple with fruiting phenology that overlaps with sympatric hawthorn trees (see Methods), providing an opportunity to examine the impact of a rare host fruit on *R. pomonella* life history timing and cold tolerance.

In this study, we examined cold tolerance and post-chill development (eclosion timing) in populations of *R. basiola* and *R. pomonella* across different elevations and host fruits in Colorado, USA. We predicted crabapple- and hawthorn-infesting *R. pomonella* would differ in cold tolerance and post-chill eclosion timing due to impacts of host fruit, either due to diet or evolved differences between populations. We predicted that *R. basiola* would be freeze-avoidant (similar to *R. pomonella*) and populations from high elevations would be more cold-tolerant and those from lower elevations. Surprisingly, we found limited impact of elevation or host fruit on cold tolerance and post-chill eclosion timing of either species, suggesting cold tolerance is relatively invariant within *R. pomonella* and *R. basiola*.

## Methods

### Insect collection

We collected host fruits of *R. basiola*, *R. pomonella*, and an unidentified species of *Rhagoletis* (subsequently identified as *R. pomonella*; see Results) in September 2018 and 2019 in and around Denver, Colorado (Table 1). To compare *R. basiola* cold tolerance at two different elevations, we obtained infested rosehips (*Rosa* sp.) from 1630 m and 2900 m above sea level (Table 1). To compare *R. pomonella* cold tolerance and eclosion across different host fruits, we collected infested hawthorn fruits (*Crataegus* sp.) and Dolgo crabapples (*Malus* ‘Dolgo’) from a very narrow geographic range in the Denver city limits. The Forest Parkway crabapples were collected earlier (early September) in the fall than the other *R. pomonella* populations (late September; Table 1). We transported all fruits to the University of Colorado, Denver within 24 hours of collection, stored the fruits at c. 22°C, and collected *Rhagoletis* pupae from these fruits over several weeks using previously described methods (McIntyre et al., 2023; Toxopeus et al., 2021). Newly formed pupae were placed in petri dishes and incubated at 21°C, 14:10 L:D for 10 days. They were then chilled at 4°C in constant darkness for up to 21 weeks until the start of the experiments described below.

**Table 1.** Collection locations of *Rhagoletis basiola* and *Rhagoletis pomonella* for this study.

| Location name | Species | Host fruit | Collection year | Location coordinates<sup>a</sup> | Location elevation (m a.s.l.)<sup>b</sup> |
| --- | --- | --- | --- | --- | --- |
| Boulder Creek | *R. basiola* | Rosehip<br>(*Rosa* sp.) | 2018 | 40.006,<br>-105.350 | 1660 |
| Chatfield Farms | *R. basiola* | Rosehip<br>(*Rosa* sp.) | 2018 | 39.577,<br>-105.096 | 1630 |
| Mountain Research Station | *R. basiola* | Rosehip<br>(*Rosa* sp.) | 2018 | 40.032,<br>-105.534 | 2900 |
| Cheesman Park | *R. pomonella* | Hawthorn<br>(*Crataegus* sp.) | 2018 | 39.733,<br>-104.966 | 1610 |
| East High | *R. pomonella* | Hawthorn<br>(*Crataegus* sp.) | 2019 | 39.742,<br>-104.956 | 1610 |
| Forest Parkway | *R. pomonella* | Hawthorn<br>(*Crataegus* sp.) | 2019 | 39.746,<br>-104.927 | 1610 |
| Forest Parkway | *R. pomonella* | Dolgo crabapple<br>(*Malus* sp.) | 2019 | 39.746,<br>-104.927 | 1610 |
| Congress Park | *R. pomonella* | Dolgo crabapple<br>(*Malus* sp.) | 2019 | 39.730,<br>-104.958 | 1610 |

<sup>a</sup>World Geodetic System WGS84 system used for coordinates; <sup>b</sup> a.s.l., above sea level

### Species identification

Dolgo crabapples are a cultivar introduced to North America as ornamental plants and derived from the Asian crab apple *Malus baccata*, a species that is relatively closely related to and interbreeds with domesticated apples (*Malus domestica*), and were introduced to North America as ornamental plants (Cornille et al., 2014; Jefferson, 1970). Given that domesticated apples are a common host fruit for *R. pomonella*, it was probable that the Dolgo crabapples were also infested with *R. pomonella*. However, reports of *Rhagoletis* spp. in or near crabapples are rare (O’Kane, 1914; St. Jean et al., 2013; Yee, 2008; Yee and Klaus, 2015, 2013). *Rhagoletis pomonella* has been reported at very low levels in residential, domesticated apple trees in Colorado prior to the study (Hood et al., 2014), but no study has detected *Rhagoletis* in crabapple in Denver, Colorado before.

We used a combination of morphological and genetic methods to confirm the species of *Rhagoletis* infesting the two sites with Dolgo crabapples. We reared some of the crabapple flies to adulthood and determined that wing morphology (see Results for example photos) was consistent with *R. pomonella*, *Rhagoletis zephyria*, and *Rhagoletis mendax* (Carroll et al., 2002). Based on the geographic location of our samples, *R. mendax* was an unlikely candidate. To narrow down the species of *Rhagoletis* infesting our Dolgo crabapples, we used a genetic assay designed to distinguish *R. pomonella* from *R. zephryia* using a single set of primers that amplifies a region of non-transcribed spacer (NTS) in the rDNA cistron, which happens to be longer in *R. pomonella* (840 bp) than in *R. zephyria* (790 bp) (Smith et al., 2022).

To conduct the genetic assay, we first extracted DNA from crabapple and hawthorn pupae using the PureGene DNA Extraction kit (Qiagen, Toronto ON, Canada) according to the manufacturer’s instructions and scaled down so that each individual was homogenized in 100 µl of cell lysis solution. We then amplified the NTS regions via PCR using 1 µL template DNA, 10 µM each PMZrDNA primers (Smith et al., 2022), and DreamTaq Polymerase (ThermoFisher Scientific, Toronto ON, Canada) in 25 µL reactions. We used the same thermocycler conditions as Smith et al. (2022): an initial denaturation step of 95°C for 3 min; 35 cycles of denaturation at 95°C for 30 s, annealing at 58°C for 30 s, and extension at 72°C for 1 min, followed by a final extension at 72°C for 5 min. We conducted agarose gel electrophoresis to determine amplicon size and Sanger Sequencing at The Centre for Applied Genomics (Sick Kids Hospital, Toronto, ON, Canada) to determine the expected target had indeed been amplified.

### Cold tolerance assays

We determined the cold tolerance strategy of four groups of *Rhagoletis* pupae (Table 1): *R. basiola* collected in 2018 from Boulder Creek, *R. pomonella* collected in 2018 and 2019 from hawthorn in Denver, and *R. pomonella* collected in 2019 from crabapple in Denver (*N* = 18 – 20 per group). This experiment was conducted on pupae that had already been chilled at 4°C for 20-21 weeks. We used previously described methods (McIntyre et al., 2023; Sinclair et al., 2015; Toxopeus et al., 2021) to cool pupae at −0.25°C/min from 4°C to a subzero temperature at which half of the individuals in the group remained supercooled and the other half froze. Ice formation is exothermic and causes a clear increase in insect body temperature at the SCP (Sinclair et al., 2015). Freezing was therefore detected through constant (once per second) temperature monitoring of each pupa with Type T thermocouples interfaced with PicoLog v6 via TC-08 units. We then transferred pupae to 21°C for 7 days to recover. We assessed survival 7 days post-cold treatment using stop-flow respirometry at 21°C (Ragland et al., 2009; Toxopeus et al., 2021). Following respirometry, we removed puparia to confirm mortality, checking for visual signs of decay or decomposition. Groups were classified as freeze-tolerant if at least 75% of them survived freezing of their body fluids, freeze-avoidant if at least 75% of them survived supercooling but not freezing, and chill-susceptible more than 75% of them did not survive supercooling or freezing (Sinclair et al., 2015).

To confirm the initial results that all groups were freeze-avoidant, we did a longer exposure to a low temperature in two of the populations described above. Groups (*N* = 20) of *R. basiola* (2018 Boulder Creek) and *R. pomonella* (2018 hawthorn) were cooled using similar methods to the cold tolerance strategy experiment. We conducted this experiment after the pupae had already been chilled at 4°C for 20-21 weeks. We cooled these groups from 4°C to −19°C at - 0.25°C/min, held at −19°C for 1 hour, and then warmed to 4°C at 0.25°C/min. No individuals froze during this treatment. Pupae were then returned to 21°C, followed by respirometry 7 days post-chill to assess survival, as described above.

To compare the acute cold tolerance among populations collected from different host fruits and different elevations, we measured SCPs after 20-21 weeks of chilling at 4°C. SCP is an appropriate metric for comparing acute cold tolerance of freeze-avoidant insects because a lower SCP indicates a greater ability to supercool (and therefore survive) at subzero temperatures (Sinclair et al., 2015). To examine the impact of elevation on acute cold tolerance, we measured SCP in all groups of *R. basiola* (*N* = 18 – 20 per group), including those from lower and higher elevations (Table 1). To determine whether populations from different host fruits differed in their acute cold tolerance, we measure SCP in *R. pomonella* from hawthorn fruits (East High) and crabapples (Forest Parkway) collected in 2019 (Table 1). We cooled pupae (as described above) to a temperature at which all pupae froze, and determined the SCP values as the lowest temperature before the exotherm caused by the latent heat of crystallization (Sinclair et al., 2015).

Because *Rhagoletis* spp. overwinter in (often snow-covered) soil (Boller and Prokopy, 1976; Dean and Chapman, 1973), they do not frequently experience extreme low temperatures. We therefore examined survival following milder but longer chilling durations. We only conducted these experiments in *R. pomonella* because we had an insufficient sample size of *R. basiola*. In 2018, we exposed *R. pomonella* pupae collected from hawthorn fruits to 4°C for 14 weeks (mild overwintering). Then, groups of 9 pupae were exposed to different durations at 0°C ranging from 1 to 16 weeks. A temperature of 0°C was achieved by creating an ice slurry in a Styrofoam box stored in a walk-in 4°C incubator (Toxopeus et al., 2016). Following chilling, pupae were allowed to recover for 1 week at 21°C before assessing survival via respirometry as described above. In 2019, we exposed c. 100 *R. pomonella* pupae collected from crabapple fruits (Forest Parkway) to 4°C for 20 weeks (mild overwintering) and then −5°C for 2 weeks using the recirculating chiller described above. Once again pupae were allowed to recover for 1 week at 21°C before assessing survival via respirometry.

### Eclosion time

To examine whether ecolosion phenology of *R. pomonella* varied among host fruits, we chilled groups of *R. pomonella* pupae from four collections sites for 20 weeks at 4°C, and then tracked time for those pupae to eclose as adults at 21°C. The four collection sites included crabapple fruits (Forest Parkway, Congress Park) and hawthorns (Forest Parkway, East High; Table 1). The Forest Parkway crabapple flies were collected in early September, while the other three collections were conducted in late September. When feasible, we also tracked eclosion of pupae from our cold tolerance experiments, which gave us both a measurement of survival to the adult stage as well as time to reach the adult stage. Eclosion was checked once per week for 100 days post-chill.

### Statistical analyses

All statistical analyses were conducted in R v4.1.0 (R Core Team, 2023). To compare SCPs among *R. basiola* groups from different elevations, we fit a linear model including a factor for elevation (low and high), collection location (Boulder Creek, Chatfield Farms, or Mountain Research Station) as a random effect within elevation, and a continuous covariate for mass. To compare SCPs among *R. pomonella* from hawthorn and apple fruits, we fit a linear model including a factor for host fruit type (hawthorn or apple) and a continuous covariate for mass. To compare the distribution of eclosion time among *R. pomonella* populations, we performed pairwise, nonparametric Kolmogorov-Smirnov (K-S) tests. All analysis code is available at https://github.com/jtoxopeus/Rbasiola-Rpomonella-coldtolerance.

## Results

### Rhagoletis pomonella infest ‘Dolgo’ crabapples

The wing morphology of the crabapple fly (Figure 1A) and hawthorn fly (Figure 1B) closely matched each other, and known *R. pomonella, R. mendax* and *R. zephyria* wing samples (Carroll et al., 2002). The genetic assay amplified regions of NTS for the crabapple and hawthorn PCR products, which both had the expected fragment size for the species *R. pomonella* (c. 840 bp; Figure 1C), rather than *R. zephyria* (790 bp). Sanger sequencing confirmed that our sequences have the diagnostic indel of *R. pomonella*, not *R. zephyria* (Figure S1, Smith et al. 2022). Taken together, the genetic assay and wing morphology data strongly suggest that Denver Dolgo crabapples were infested by *R. pomonella*.

**Figure 1.**
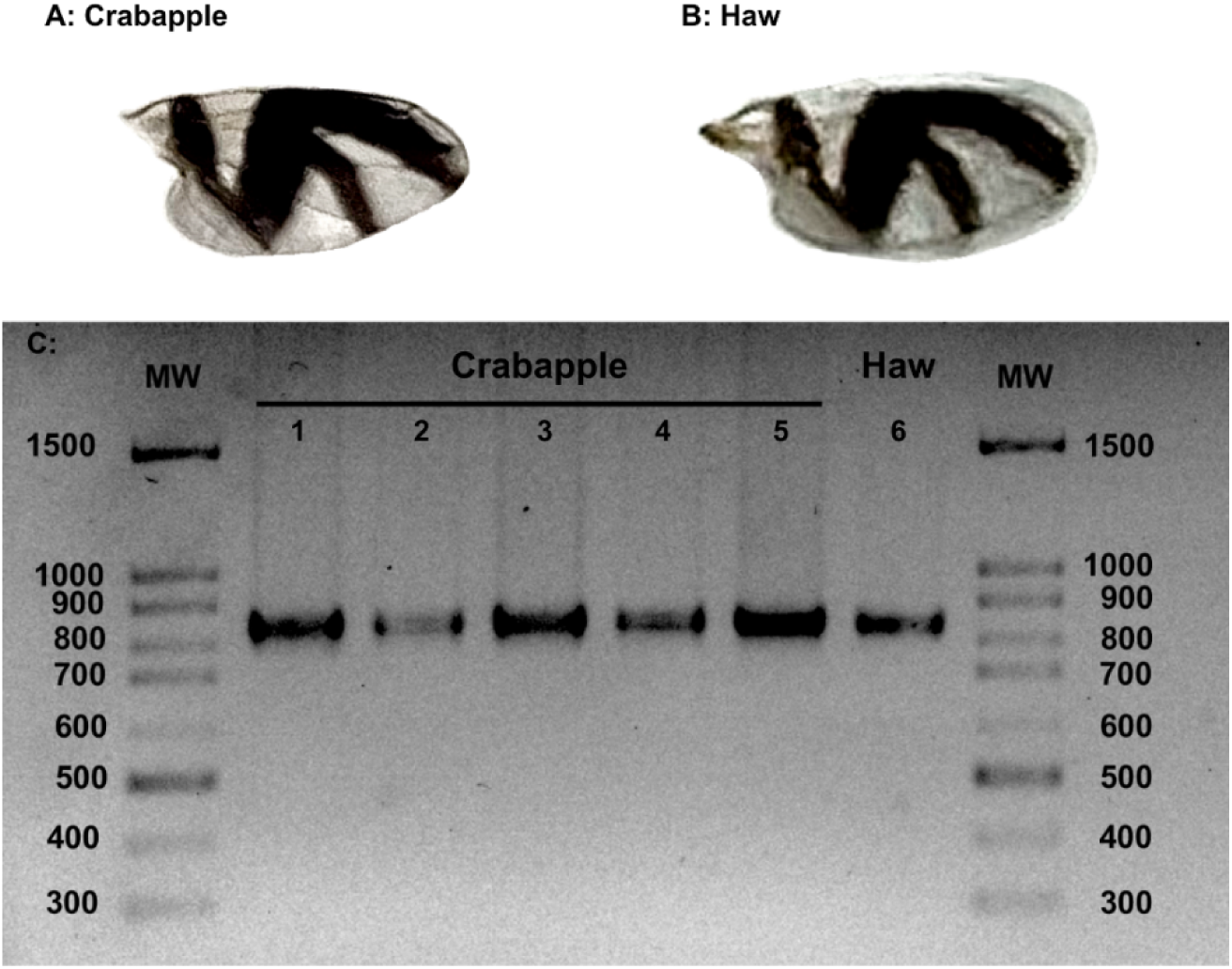
Morphological and genetic evidence for *R. pomonella* infestation of Dolgo crabapple in Denver, CO. Representative wing morphology of flies collected from **(A)** Dolgo crabapple and **(B)** hawthorn are similar. **(C)** PCR products of NTS from crabapple and hawthorn flies are the fragment length size for *R. pomonella* (∼840 bp; Smith et al., 2022). Lanes: MW, 100 bp FroggaBio ladder; lanes 1–5, crabapple flies and lane 6; hawthorn fly. Agarose gel (1.5%) was run for 75 min at 150 V and stained with RedSafe.

### Rhagoletis basiola acute cold tolerance does not vary with elevation

*Rhagoletis basiola* from both high (2900 m above sea level) and lower (1630 – 1660 m above sea level) elevations had similar tolerance to acute cold exposure. Similar to *R. pomonella*, *R. basiola* were freeze-avoidant (Table 2). Based on respirometry measurements, all *R. basiola* (Boulder Creek collection) survived supercooling at −19°C for 1 h (Table 3). The majority (13 out of 18) of these pupae eclosed as adults, and the remaining pupae appeared alive but undeveloped (i.e., likely still in diapause, cf. (Toxopeus et al., 2021) at the end of the experiment. The *R. basiola* collected from a high elevation trended towards having a lower mass (6.08 ± 0.19 mg) than those from lower elevations (13.03 ± 0.30 mg). Despite this, there was no difference in SCP between low and high elevation populations (Figure 2; Mass: *P* = 0.104; Elevation: *P* = 0.292, Population: *P* = 0.666). Given that low and high elevation *R. basiola* can supercool to similarly low temperatures, and they are freeze-avoidant, both populations have similar tolerance to acute low temperature exposures.

**Figure 2.**
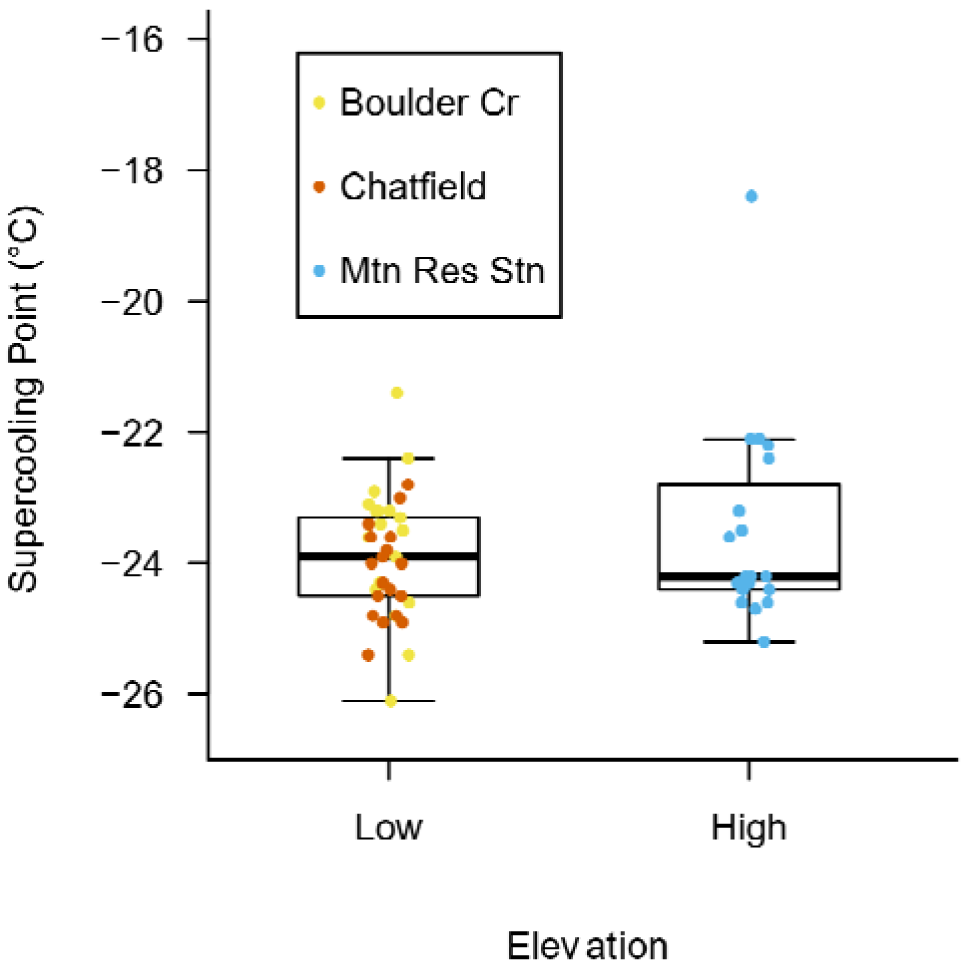
Supercooling point (SCP) temperatures of *Rhagoletis basiola* do not differ between low and high elevation populations. Each circular point represents the SCP from one individual. Different colours indicate different collection locations: yellow, Boulder Creek (1660 m above sea level); orange, Chatfield Farms (1630 m); blue, N-line; grey, Mountain Research Station (2900 m). Pupae were exposed to 4°C for 20 weeks prior to measuring SCP.

**Table 2.** Cold tolerance strategy was similar among collections of *Rhagoletis* spp. from different host fruits and across years. Survival post-cold stress was determined by respirometry and visual observations of decay. Eclosion (completion of development) was also tracked for the 2018 populations, as indicated in parentheses.

| Species | Population<br>(Fruit, Year) | Mean mass<br>± s.e. (mg) | N frozen<br>survived/<br>N frozen | N supercooled<br>survived/<br>N supercooled | Cold<br>tolerance<br>strategy |
| --- | --- | --- | --- | --- | --- |
| *Rhagoletis*<br>*basiola* | Rosehip,<br>2018<sup>a</sup> | 12.54 ± 0.31 | 0/10 | 8/9<br>(8/9 eclosed) | Freeze-avoidant |
| *Rhagoletis*<br>*pomonella* | Hawthorn,<br>2018<sup>b</sup> | 7.94 ± 0.38 | 0/11 | 7/7<br>(6/7 eclosed) | Freeze-avoidant |
| *Rhagoletis*<br>*pomonella* | Hawthorn,<br>2019<sup>c</sup> | 8.05 ± 0.27 | 0/8 | 9/11 | Freeze-avoidant |
| *Rhagoletis*<br>*pomonella* | Crabapple,<br>2019<sup>d</sup> | 6.68 ± 0.30 | 0/7 | 9/11 | Freeze-avoidant |

<sup>a</sup>Boulder Creek, <sup>b</sup>Cheesman Park, <sup>c</sup>East High, <sup>d</sup>Forest Parkway collections

**Table 3.**
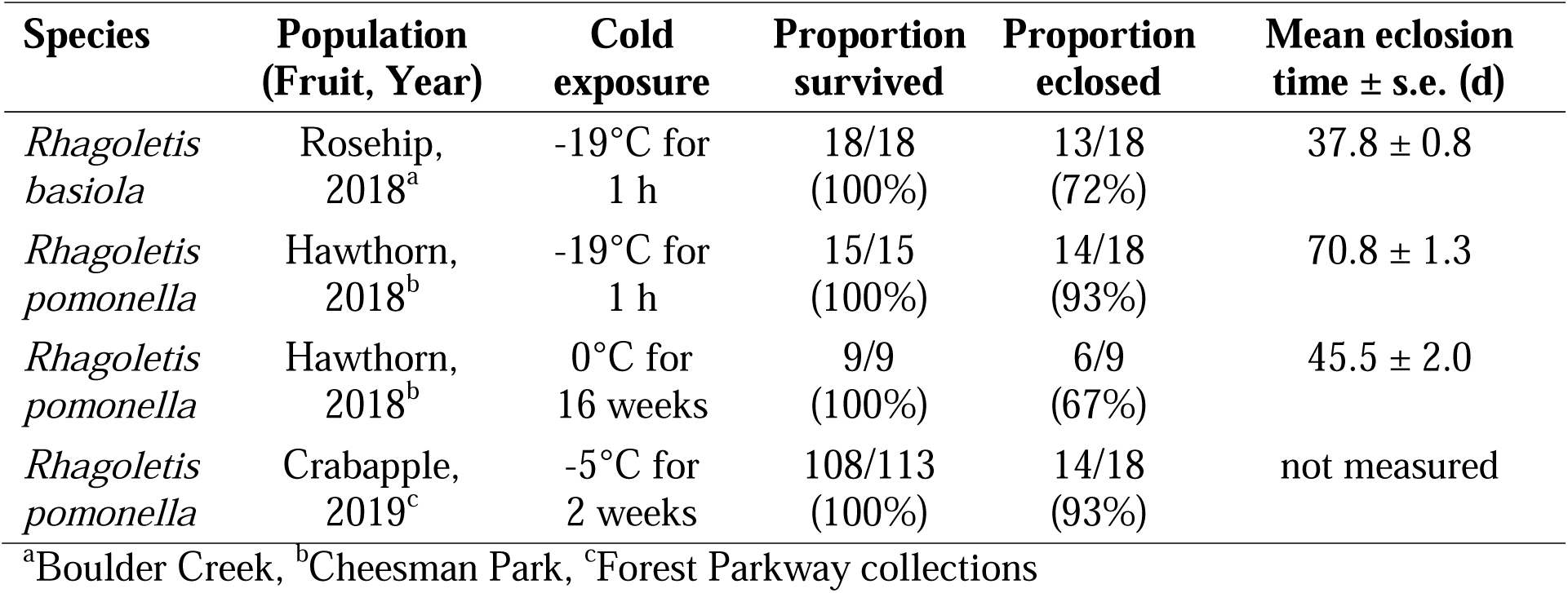
*Rhagoletis* spp. survival is high following acute exposure to −19°C or longer exposure to 0°C and −5°C. Pupae were chilled for 14-20 weeks at 4°C prior to cold exposure. Survival post-cold stress was determined by respirometry and eclosion tracking (completion of development) at 21°C. Full results for shorter exposures to 0°C are in Table S2.

| Species | Population<br>(Fruit, Year) | Cold<br>exposure | Proportion<br>survived | Proportion<br>eclosed | Mean eclosion<br>time ± s.e. (d) |
| --- | --- | --- | --- | --- | --- |
| *Rhagoletis*<br>*basiola* | Rosehip,<br>2018<sup>a</sup> | -19°C for<br>1 h | 18/18<br>(100%) | 13/18<br>(72%) | 37.8 ± 0.8 |
| *Rhagoletis*<br>*pomonella* | Hawthorn,<br>2018<sup>b</sup> | -19°C for<br>1 h | 15/15<br>(100%) | 14/18<br>(93%) | 70.8 ± 1.3 |
| *Rhagoletis*<br>*pomonella* | Hawthorn,<br>2018<sup>b</sup> | 0°C for<br>16 weeks | 9/9<br>(100%) | 6/9<br>(67%) | 45.5 ± 2.0 |
| *Rhagoletis*<br>*pomonella* | Crabapple,<br>2019<sup>c</sup> | -5°C for<br>2 weeks | 108/113<br>(100%) | 14/18<br>(93%) | not measured |

<sup>a</sup>Boulder Creek, <sup>b</sup>Cheesman Park, <sup>c</sup>Forest Parkway collections

### Rhagoletis pomonella acute cold tolerance does not vary with host fruit

*Rhagoletis pomonella* from both crabapple and hawthorn fruits had similar tolerance of acute exposures to low temperatures. Pupae collected from both fruits were freeze-avoidant (Table 2); they survived cooling to low temperatures as long as internal ice formation did not occur. If held in a supercooled state for 1 h at a low temperature above the SCP (−19°C), all *R. pomonella* from hawthorn fruits (2018 collection) were alive 1 week post-chill, and almost all (14 out of 15) of these individuals eclosed as adults (Table 3). This confirms that acute exposures to extreme low temperatures do not negatively impact post-chilling development. All *R. pomonella* pupae had a low SCP regardless of host fruit (Figure 2), and mass did not substantially impact SCP (Mass: *P* = 0.823; Fruit: *P* = 0.328). This similar supercooling ability suggests that *R. pomonella* from both hawthorns and crabapples can remain unfrozen (and therefore survive) at similarly low temperatures.

### Rhagoletis pomonella chronic cold tolerance is high

*Rhagoletis pomonella* typically have good tolerance to prolonged exposure to mild low temperatures, similar to their overwintering microhabitat in the soil. We did not have the sample sizes to directly compare prolonged chill tolerance among all of our *R. pomonella* collections. However, we can report that *R. pomonella* pupae collected from crabapples (2019 collection) had 95% survival after −5°C for 2 weeks (longer durations not tested; Table 3). In addition, almost every duration of exposure to 0°C (1 – 16 weeks) resulted in 100% survival of pupae from hawthorns (2018 collection; Table 3), except the 14 week treatment (78% survival; Table S1). These temperatures are both lower than the typical lab ‘winter’ temperature (4°C) used in most *R. pomonella* work (Calvert et al., 2022; Dambroski and Feder, 2007; Dowle et al., 2020; Feder et al., 1997; McIntyre et al., 2023; Toxopeus et al., 2021), and our results indicate good chronic chill tolerance in *R. pomonella* collected from both host fruits.

### Eclosion time varies among Rhagoletis pomonella populations

Following chilling for 20 weeks at 4°C, *R. pomonella* eclosed as adults within 100 days at 21°C (Figure 4). Crabapple flies from Forest Parkway (pupae collected in early September) eclosed about 10 days earlier than both hawthorn flies (Forest Parkway and East High) and crabapple flies (Congress Park) from other locations (pupae collected in late September; Figure 4, Table S2). However, hawthorn and crapabble flies collected at a similar time of year (late September) had similar eclosion patterns under our controlled laboratory settings (Figure 4; Table S2). This suggests that pupae that form earlier in the autumn will eclose quicker after a simulated winter than pupae collected later in the autumn and overwintered under identical conditions.

**Figure 3.**
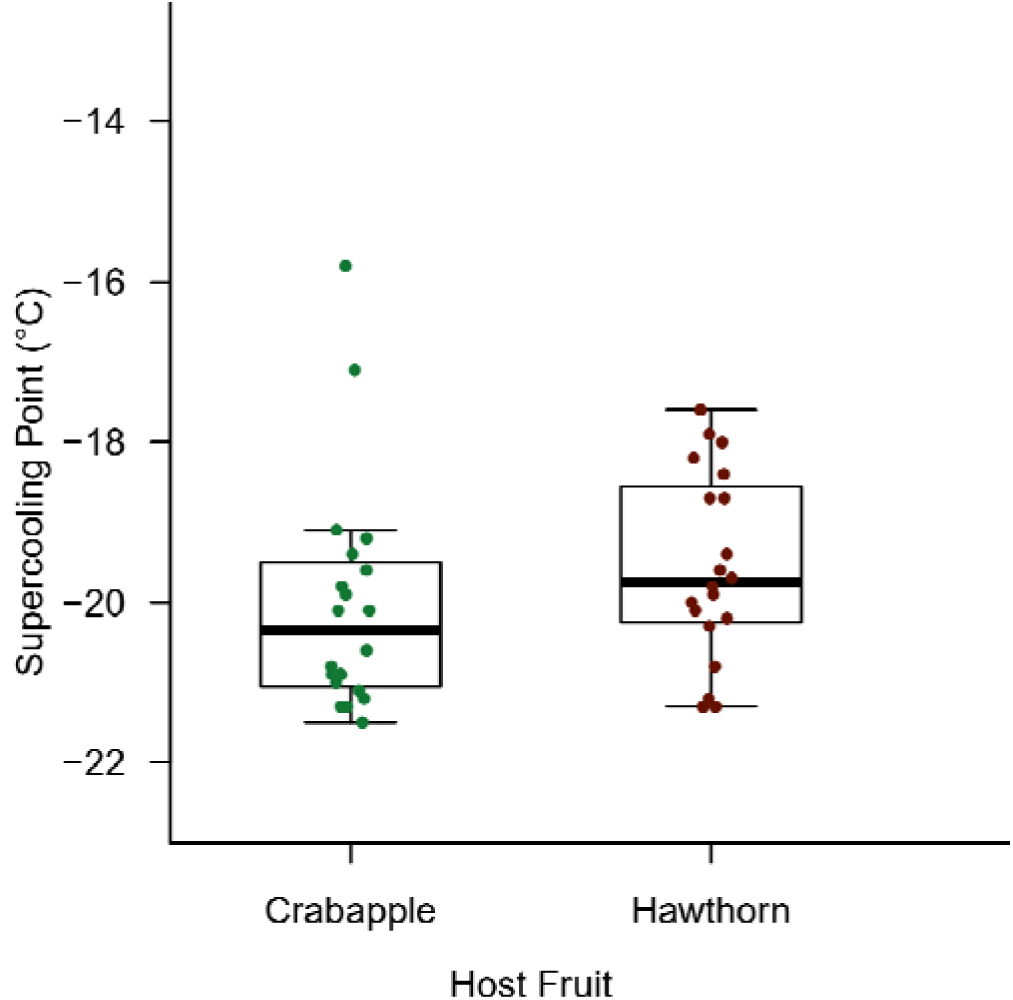
Supercooling point (SCP) temperatures of *Rhagoletis pomonella* do not differ between populations collected from crabapple and hawthorn host fruits. Each circular point represents the SCP from one individual. Crabapples were collected from Forest Parkway and hawthorns were collected from East High within the same two weeks in 2019. Pupae were exposed to 4°C for 20 weeks prior to measuring SCP.

**Figure 4.**
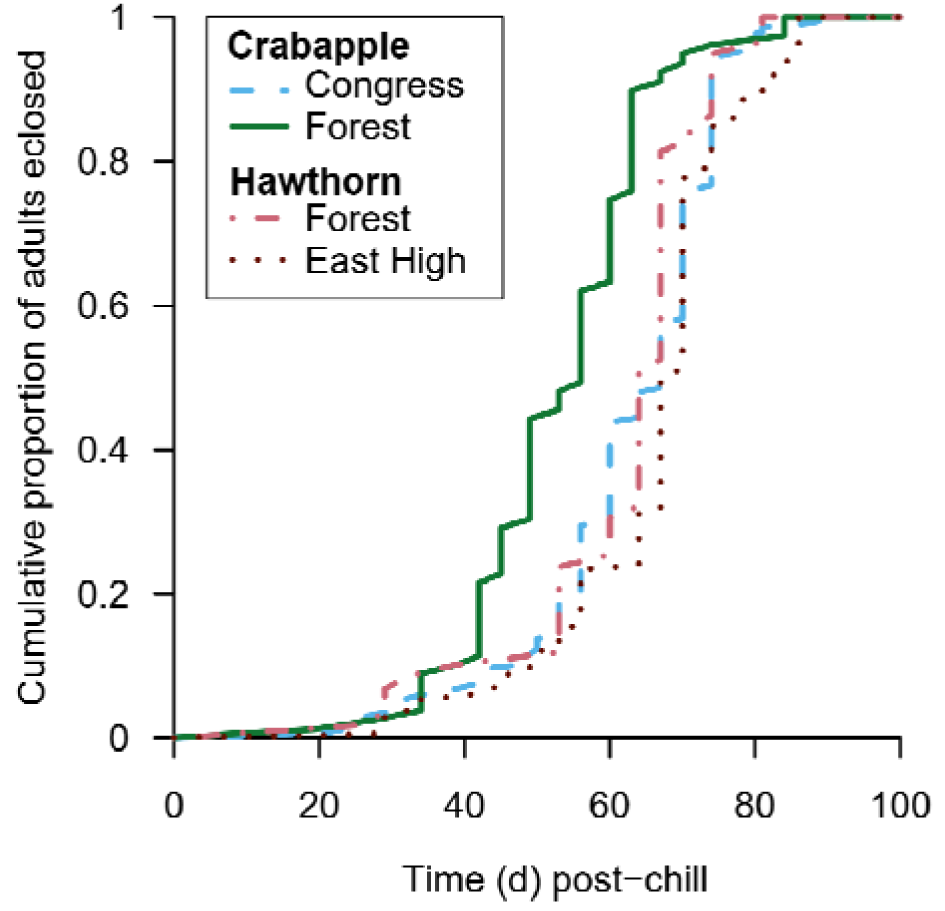
Cumulative proportion of *Rhagoletis pomonella* flies that eclosed post-chill differed among populations. Forest Parkway populations of crabapple flies were collected in early September, while the other three populations were collected in late September and early October. All pairwise comparisons of eclosion distributions between populations were significantly different, except for the Congress Park crabapple and East High hawthorn populations (Table S2). Eclosion was tracked within 100 days at 21°C after pupae were chilled a 4°C for 20 weeks.

## Discussion

In this study, we documented the presence of *R. pomonella* in an unusual host fruit (Dolgo crabapple), and characterized cold tolerance and post-chill development in previously unstudied populations of both *R. basiola* and *R. pomonella* in Colorado, USA. Contrary to our predictions, the acute cold tolerance of *R. basiola* and *R. pomonella* did not vary with elevation or host fruit, respectively. All populations were freeze-avoidant, survived acute exposures to low temperatures well, and exhibited good chronic cold tolerance. Despite the lack of differences among populations, this study lays the groundwork for continuing to examine the overwintering and post-winter survival in two interesting biological systems.

Our work on the multiple populations of *R. basiola* is the first to characterize the cold tolerance of this species. We expected cold tolerance to be stronger at high elevation, similar to other species (Dennis et al., 2015; Oyen et al., 2016; Tonione et al., 2020; Vrba et al., 2017). In addition, we expected individuals with a lower mass (high elevation *R. basiola*) to have lower SCPs (cf. (Sinclair et al., 2015), but detected no such difference. Two clear questions arise from these results. Firstly, what mechanisms underlie such a uniformity in SCP among these *R. basiola* populations? It may be that the SCP is not a particularly relevant metric of cold tolerance in this species if soil temperatures at high elevations stay relatively mild, which is likely if these high elevation sites have an insulating layer of snow cover (cf. (Roberts et al., 2021). Studies that attempt to induce plasticity (changes) in *R. basiola* cold tolerance (cf. (Butterson et al., 2021) and compare cryoprotectants and antifreeze proteins among populations of *R. basiola* would help further understand the mechanisms affecting SCP and its relevance in this species. Secondly, although acute cold tolerance did not differ within *R. basiola*, how do other metrics of cold tolerance (e.g., chronic cold tolerance) differ? Body size has implications for energy reserve allocation and survival of cold stress (Colinet et al., 2006; Renault et al., 2003), so high elevation (smaller) *R. basiola* may have weaker chronic cold tolerance than their lower elevation counterparts. Given the broad geographic distribution of *R. basiola*, it would also be worth studying whether cold tolerance differs on a larger geographic scale, similar to other insects (Addo-Bediako et al., 2000; Alfaro-Tapia et al., 2021; Sunday et al., 2011).

The *R. pomonella* crabapple and hawthorn system we examined in this study has several differences to the classic domesticated apple vs. hawthorn systems that have previously been used to study diapause and post-winter eclosion (Calvert et al., 2022; Dambroski and Feder, 2007; Dowle et al., 2020; Feder et al., 1997). First, *Rhagoletis* spp. are rarely found in or near crabapples (St. Jean et al., 2013; Yee, 2008; Yee and Klaus, 2015, 2013), and crabapples are generally considered an unsuitable host for *R. pomonella* given their acidity, late fruiting phenology, and phenolic content (Bush et al., 1989); but see (Neilson, 1967). There is at least one record of infestation in a Dolgo variety apple (O’Kane, 1914) in the eastern US, but the author and original observer attribute this to the proximity of the tree to a very heavily infested apple orchard. Not only did we detect *R. pomonella* in Dolgo crabapples (an introduced ornamental from Russia; (Jefferson, 1970) in Denver, but we detected them in similar abundance and at similar times to the *R. pomonella* in nearby hawthorn trees. In addition, when we collected host fruits at similar times (late September), the post-chill eclosion phenology did not differ between crabapple and hawthorn flies. This is in contrast to *R. pomonella* collected from domesticated apples, which tend to eclose faster post-chilling than those collected from hawthorn fruits when overwintered under common conditions (Feder et al., 1993; Filchak et al., 2000; Ragland et al., 2017; Smith, 1988). Further research is required to determine whether these crabapple flies are a distinct population from the nearby hawthorn flies – similar to domesticated apple vs. hawthorn flies; (Dowle et al., 2020; Filchak et al., 2000; Ragland et al., 2017) – or whether they form a single population using two host fruits. The latter would provide an interesting study system to examine the impact of host fruit on biological processes (e.g., diapause) without the impact of substantial genetic (evolved) differences between flies using those host fruits.

In conclusion, we have shown a relatively conserved level of cold tolerance among populations within each species in this study, *R. basiola* and *R. pomonella*. Future work that examines cold tolerance mechanisms in these species may further increase our understanding of the (lack of) variation in cold tolerance in *Rhagoletis* spp. In addition, our documentation of *R. pomonella* in Dolgo crabapples in close proximity to hawthorn trees provides an additional system to examine how host fruits impact the biology of *R. pomonella*.

## Data Availability Statement

All data and analysis code are available at https://github.com/jtoxopeus/Rbasiola-Rpomonella-coldtolerance.

## Ethics Statement

Ethics Approval was not necessary for this study.

## Acknowledgements

Thank you to the following, who helped with collecting and processing *Rhagoletis* samples: L. Andaloori, M. Calvert, E. Kelso, K. Marshall, M. Marquis, I. Ragland, L. Ragland, M. Sanaei, I. Sower, J. Tucker, B. Vestby

## Author Contributions

All authors contributed to revising the manuscript. In addition, KL: data collection and analysis, manuscript original draft; MM, PB, and LG: data collection; GJR: conceptualization; JT: conceptualization, data collection and analysis.

## References

Addo-Bediako, A., Chown, S.L., Gaston, K.J., 2000. Thermal tolerance, climatic variability and latitude. Proc. R. Soc. B Biol. Sci. 267, 739–745. 10.1098/rspb.2000.1065

Alfaro-Tapia, A., Alvarez-Baca, J., Tougeron, K., Lavandero, B., Le Lann, C., van Baaren, J., 2021. Overwintering strategies and life-history traits of different populations of *Aphidius platensis* along a latitudinal gradient in Chile. Entomol. Gen. 122, 127–145. 10.1127/entomologia/2021/1186

Averill, A.L., Prokopy, R.J., 1981. Oviposition deterring fruit marking pheromone in *Rhagoletis basiola*. Fla. Entomol. 64, 222–226. 10.2307/3494573

Berlocher, S.H., Dixon, P.L., 2004. Occurrence of Rhagoletis species in Newfoundland. Entomol. Exp. Appl. 113, 45–52. 10.1111/j.0013-8703.2004.00204.x

Boller, E.F., Prokopy, R.J., 1976. Bionomics and management of *Rhagoletis*. Annu. Rev. Entomol. 21, 223–246. 10.1146/annurev.en.21.010176.001255

Bush, G.L., Feder, J.L., Berlocher, S.H., McPheron, B., Smith, Dc., Chilcote, C., 1989. Sympatric origins of *R. pomonella*. Nature 339, 346.

Butterson, S., Roe, A.D., Marshall, K.E., 2021. Plasticity of cold hardiness in the eastern spruce budworm, *Choristoneura fumiferana*. Comp. Biochem. Physiol. A. Mol. Integr. Physiol. 259, 110998. 10.1016/j.cbpa.2021.110998

Calvert, M.B., Doellman, M.M., Feder, J.L., Hood, G.R., Meyers, P., Egan, S.P., Powell, T.H.Q., Glover, M.M., Tait, C., Schuler, H., Berlocher, S.H., Smith, J.J., Nosil, P., Hahn, D.A., Ragland, G.J., 2022. Genomically correlated trait combinations and antagonistic selection contributing to counterintuitive genetic patterns of adaptive diapause divergence in *Rhagoletis* flies. J. Evol. Biol. 35, 146–163. 10.1111/jeb.13952

Canadian Food Inspection Agency, 2012. Rhagoletis pomonella (Apple maggot) - Fact sheet [WWW Document]. URL https://inspection.canada.ca/plant-health/invasive-species/insects/apple-maggot/fact-sheet/eng/1330366145611/1330366375524 (accessed 7.20.23).

Carroll, L.E., White, I.M., Freidburg, A., Norbomm, A.L., Dallwitz, M.J., Thompson, F.C., 2002. Pest fruit flies of the world [WWW Document]. URL https://www.delta-intkey.com/ffa/index.htm (accessed 12.4.23).

Colinet, H., Hance, T., Vernon, P., 2006. Water relations, fat reserves, survival, and longevity of a cold-exposed parasitic wasp *Aphidius colemani* (Hymenoptera: Aphidiinae). Environ. Entomol. 35, 228–236. 10.1603/0046-225X-35.2.228

Cornille, A., Giraud, T., Smulders, M.J.M., Roldán-Ruiz, I., Gladieux, P., 2014. The domestication and evolutionary ecology of apples. Trends Genet. 30, 57–65. 10.1016/j.tig.2013.10.002

Dambroski, H.R., Feder, J.L., 2007. Host plant and latitude-related diapause variation in *Rhagoletis pomonella*: a test for multifaceted life history adaptation on different stages of diapause development. J. Evol. Biol. 20, 2101–2112. 10.1111/j.1420-9101.2007.01435.x

Dean, R.L., Chapman, P.J., 1973. Bionomics of the apple maggot in eastern New York. Search Agric. 3, 1–64.

Denlinger, D.L., 2022. Insect Diapause. Cambridge University Press, Cambridge. 10.1017/9781108609364

Dennis, A.B., Dunning, L.T., Sinclair, B.J., Buckley, T.R., 2015. Parallel molecular routes to cold adaptation in eight genera of New Zealand stick insects. Sci. Rep. 5, 13965. 10.1038/srep13965

Doellman, M.M., Hood, G.R., Gersfeld, J., Driscoe, A., Xu, C.C.Y., Sheehy, R.N., Holmes, N., Yee, W.L., Feder, J.L., 2020. Identifying diagnostic genetic markers for a cryptic invasive agricultural pest: A test case using the apple maggot fly (Diptera: Tephritidae). Ann. Entomol. Soc. Am. 113, 246–256. 10.1093/aesa/saz069

Dowle, E.J., Powell, T.H.Q., Doellman, M.M., Meyers, P.J., Calvert, M.B., Walden, K.K.O., Robertson, H.M., Berlocher, S.H., Feder, J.L., Hahn, D.A., Ragland, G.J., 2020. Genome-wide variation and transcriptional changes in diverse developmental processes underlie the rapid evolution of seasonal adaptation. Proc. Natl. Acad. Sci. 117, 23960–23969. 10.1073/pnas.2002357117

Feder, J.L., Hunt, T.A., Bush, L., 1993. The effects of climate, host plant phenology and host fidelity on the genetics of apple and hawthorn infesting races of *Rhagoletis pomonella*. Entomol. Exp. Appl. 69, 117–135. 10.1111/j.1570-7458.1993.tb01735.x

Feder, J.L., Stolz, U., Lewis, K.M., Perry, W., Roethele, J.B., Rogers, A., 1997. The effects of winter length on the genetics of apple and hawthorn races of *Rhagoletis pomonella* (Diptera: Tephritidae). Evolution 51, 1862–1876. 10.1111/j.1558-5646.1997.tb05109.x

Filchak, K.E., Roethele, J.B., Feder, J.L., 2000. Natural selection and sympatric divergence in the apple maggot *Rhagoletis pomonella*. Nature 407, 739–742.

Hahn, D.A., Denlinger, D.L., 2011. Energetics of insect diapause. Annu. Rev. Entomol. 56, 103–121. 10.1146/annurev-ento-112408-085436

Hand, S.C., Denlinger, D.L., Podrabsky, J.E., Roy, R., 2016. Mechanisms of animal diapause: recent developments from nematodes, crustaceans, insects, and fish. Am. J. Physiol. 310, R1193–R1211. 10.1152/ajpregu.00250.2015

Hodek, I., 1996. Diapause development, diapause termination and the end of diapause. Eur. J. Entomol. 93, 475–487.

Hoffmeister, T.S., Lachlan, R.F., Roitberg, B.D., 1999. Do larger fruits provide a partial refuge for rose-hip flies against parasitoids? J. Insect Behav. 12, 451–460. 10.1023/A:1020906521745

Hoffmeister, T.S., Roitberg, B.D., Lalonde, R.G., 2000. Catching Ariadne by her thread: how a parasitoid exploits the herbivore’s marking trails to locate its host. Entomol. Exp. Appl. 95, 77–85. 10.1046/j.1570-7458.2000.00644.x

Hood, G.R., Glover, M., Tait, C., Yee, W.L., Feder, J.L., 2014. Detection of an apple-infesting population of *Rhagoletis pomonella* (Walsh 1867) (Diptera: Tephritidae) in the state of Colorado, USA. Pan-Pac. Entomol. 90, 4–10. 10.3956/2014-90.1.4

Hood, G.R., Yee, W., Goughnour, R.B., Sim, S.B., Egan, S.P., Arcella, T., Saint-Jean, G., Powell, T.H.Q., Xu, C.C.Y., Feder, J.L., 2013. The geographic distribution of *Rhagoletis pomonella* (Diptera: Tephritidae) in the western United States: introduced species or native population? Ann. Entomol. Soc. Am. 106, 59–65. 10.1603/AN12074

Jefferson, R.M., 1970. History, Progeny, and Locations of Crabapples of Documented Authentic Origin. Agricultural Research Service, U. S. Department of Agriculture.

Koštál, V., 2006. Eco-physiological phases of insect diapause. J. Insect Physiol. 52, 113–127. 10.1016/j.jinsphys.2005.09.008

Koštál, V., Korbelová, J., Poupardin, R., Moos, M., Šimek, P., 2016. Arginine and proline applied as food additives stimulate high freeze tolerance in larvae of *Drosophila melanogaster*. J. Exp. Biol. 219, 2358–2367. 10.1242/jeb.142158

Koštál, V., Zahradníčková, H., Šimek, P., 2011. Hyperprolinemic larvae of the drosophilid fly, Chymomyza costata, survive cryopreservation in liquid nitrogen. Proc. Natl. Acad. Sci. 108, 13041–13046. 10.1073/pnas.1107060108

Lee, R.E., 2010. A primer on insect cold-tolerance, in: Denlinger, D.L., Lee, R.E. (Eds.), Low Temperature Biology of Insects. Cambridge University Press, New York, pp. 3–34.

Li, Y., Zhang, L., Zhang, Q., Chen, H., Denlinger, D.L., 2014. Host diapause status and host diets augmented with cryoprotectants enhance cold hardiness in the parasitoid *Nasonia vitripennis*. J. Insect Physiol. 70, 8–14. 10.1016/j.jinsphys.2014.08.005

McIntyre, T., Andaloori, L., Hood, G.R., Feder, J.L., Hahn, D.A., Ragland, G.J., Close, Toxopeus, J., 2023. Cold tolerance and diapause within and across trophic levels: Endoparasitic wasps and their fly host have similar phenotypes. J. Insect Physiol. 146, 104501. 10.1016/j.jinsphys.2023.104501

Neilson, W.T.A., 1967. Development and mortality of the apple maggot, *Rhagoletis pomonella* in crab apples. Can. Entomol. 99, 217–219. 10.4039/Ent99217-2

O’Kane, W.C., 1914. The apple maggot. N. H. Exp. Stn. Bull. 171, 120.

Overgaard, J., MacMillan, H.A., 2017. The integrative physiology of insect chill tolerance. Annu. Rev. Physiol. 79, 187–208. 10.1146/annurev-physiol-022516-034142

Oyen, K.J., Giri, S., Dillon, M.E., 2016. Altitudinal variation in bumble bee (*Bombus*) critical thermal limits. J. Therm. Biol. 59, 52–57. 10.1016/j.jtherbio.2016.04.015

Ragland, G.J., Doellman, M.M., Meyers, P.J., Hood, G.R., Egan, S.P., Powell, T.H., Hahn, D.A., Nosil, P., Feder, J.L., 2017. A test of genomic modularity among lifeLhistory adaptations promoting speciation with gene flow. Mol. Ecol. 26, 3926–3942. 10.1111/mec.14178

Ragland, G.J., Fuller, J., Feder, J.L., Hahn, D.A., 2009. Biphasic metabolic rate trajectory of pupal diapause termination and post-diapause development in a tephritid fly. J. Insect Physiol. 55, 344–350. 10.1016/j.jinsphys.2008.12.013

Ragland, G.J., Sim, S.B., Goudarzi, S., Feder, J.L., Hahn, D.A., 2012. Environmental interactions during host race formation: host fruit environment moderates a seasonal shift in phenology in host races of *Rhagoletis pomonella*. Funct. Ecol. 26, 921–931. 10.1111/j.1365-2435.2012.01992.x

R Core Team, 2023.R: A Language and Environment for Statistical Computing.

Renault, D., Hance, T., Vannier, G., Vernon, P., 2003. Is body size an influential parameter in determining the duration of survival at low temperatures in *Alphitobius diaperinus* Panzer (Coleoptera: Tenebrionidae)? J. Zool. 259, 381–388. 10.1017/S0952836902003382

Roberts, K.T., Rank, N.E., Dahlhoff, E.P., Stillman, J.H., Williams, C.M., 2021. Snow modulates winter energy use and cold exposure across an elevation gradient in a montane ectotherm. Glob. Change Biol. 27, 6103–6116. 10.1111/gcb.15912

Roitberg, B.D., Lalonde, R.G., 1991. Host marking enhances parasitism risk for a fruit-Infesting fly *Rhagoletis basiola*. Oikos 61, 389–393. 10.2307/3545246

Rull, J., Aluja, M., Feder, J.L., Berlocher, S.H., 2009. Distribution and host range of hawthorn-infesting *Rhagoletis* (Diptera: Tephritidae) in Mexico. Ann. Entomol. Soc. Am. 99, 662– 672. 10.1603/0013-8746(2006)99[662:DAHROH]2.0.CO;2.

Sinclair, B.. J., 1999. Insect cold tolerance: how many kinds of frozen? Eur. J. Entomol. 96, 157– 164.

Sinclair, B.J., Coello Alvarado, L.E., Ferguson, L.V., 2015. An invitation to measure insect cold tolerance: Methods, approaches, and workflow. J. Therm. Biol. 53, 180–197. 10.1016/j.jtherbio.2015.11.003

Smith, D.C., 1988. Heritable divergence of *Rhagoletis pomonella* host races by seasonal asynchrony. Nature 336, 66–67. 10.1038/336066a0

Smith, J.J., Brzezinski, P., Dziedziula, J., Rosenthal, E., Klaus, M., 2022. Partial ribosomal nontranscribed spacer sequences distinguish *Rhagoletis zephyria* (Diptera: Tephritidae) from the apple maggot, R. pomonella. J. Econ. Entomol. 115, 647–661. 10.1093/jee/toab264

St. Jean, G., Egan, S.P., Yee, W.L., Feder, J.L., 2013. Genetic identification of an unknown *Rhagoletis* fruit fly (Diptera: Tephritidae) infesting Chinese crabapple: implications for apple pest management. J. Econ. Entomol. 106, 1511–1515. 10.1603/EC12449

Sunday, J.M., Bates, A.E., Dulvy, N.K., 2011. Global analysis of thermal tolerance and latitude in ectotherms. Proc. R. Soc. B Biol. Sci. 278, 1823–1830. 10.1098/rspb.2010.1295

Teets, N.M., Denlinger, D.L., 2013. Physiological mechanisms of seasonal and rapid coldLhardening in insects. Physiol. Entomol. 38, 105–116.

Tonione, M.A., Cho, S.M., Richmond, G., Irian, C., Tsutsui, N.D., 2020. Intraspecific variation in thermal acclimation and tolerance between populations of the winter ant, *Prenolepis imparis*. Ecol. Evol. 10, 4749–4761. 10.1002/ece3.6229

Toxopeus, J., Dowle, E.J., Andaloori, L., Ragland, G.J., 2023. Variation in thermal sensitivity of diapause development among individuals and over time drives life history timing patterns in an insect pest. 10.1101/2023.05.31.543112

Toxopeus, J., Gadey, L., Andaloori, L., Sanaei, M., Ragland, G.J., 2021. Costs of averting or prematurely terminating diapause associated with slow decline of metabolic rates at low temperature. Comp. Biochem. Physiol. A 255, 110920. 10.1016/j.cbpa.2021.110920

Toxopeus, J., Jakobs, R., Ferguson, L.V., Gariepy, T.D., Sinclair, B.J., 2016. Reproductive arrest and stress resistance in winter-acclimated *Drosophila suzukii*. J. Insect Physiol. 89, 37–51. 10.1016/j.jinsphys.2016.03.006

Toxopeus, J., McKinnon, A.H., Štětina, T., Turnbull, K.F., Sinclair, B.J., 2019. Laboratory acclimation to autumn-like conditions induces freeze tolerance in the spring field cricket *Gryllus veletis* (Orthoptera: Gryllidae). J. Insect Physiol. 113, 9–16. 10.1016/j.jinsphys.2018.12.007

Toxopeus, J., Sinclair, B.J., 2018. Mechanisms underlying insect freeze tolerance. Biol. Rev. 93, 1891–1914. 10.1111/brv.12425

Vrba, P., Nedvěd, O., Zahradníčková, H., Konvicka, M., 2017. More complex than expected: Cold hardiness and the concentration of cryoprotectants in overwintering larvae of five *Erebia* butterflies (Lepidoptera: Nymphalidae). Eur. J. Entomol. 114, 470–480. 10.14411/eje.2017.060

Williams, C.M., Henry, H.A., Sinclair, B.J., 2015. Cold truths: how winter drives responses of terrestrial organisms to climate change. Biol. Rev. 90, 214–35. 10.1111/brv.12105

Wilsterman, K., Ballinger, M.A., Williams, C.M., 2021. A unifying, eco-physiological framework for animal dormancy. Funct. Ecol. 35, 11–31. 10.1111/1365-2435.13718

Yee, W., Goughnour, R.B., 2006. New host records for the apple maggot, *Rhagoletis pomonella* (Walsh) (Diptera: Tephritidae), in Washington State. Pan-Pac. Entomol. 82, 54–60.

Yee, W.E., Hernández-Ortiz, V., Rull, J., Sinclair, B.J., 2013. Status of *Rhagoletis* (Diptera: Tephritidae) pests in the NAPPO Countries, NAPPO Science and Technology Documents.

Yee, W.L., 2008. Host plant use by apple maggot, western cherry fruit fly, and other *Rhagoletis* species (Diptera: Tephritidae) in central Washington state. Pan-Pac. Entomol. 84, 163–178. 10.3956/2007-48.1

Yee, W.L., Goughnour, R.B., Feder, J.L., 2021. Distinct adult eclosion traits of sibling species *Rhagoletis pomonella* and *Rhagoletis zephyria* (Diptera: Tephritidae) under laboratory conditions. Environ. Entomol. 50, 173–182. 10.1093/ee/nvaa148

Yee, W.L., Klaus, M.W., 2015. Implications of *Rhagoletis zephyria* Snow, 1894 (Diptera: Tephritidae) captures for apple maggot surveys and fly ecology in Washington State, U.S.A. Pan-Pac. Entomol. 91, 305–317. 10.3956/2015-91.4.305

Yee, W.L., Klaus, M.W., 2013. Development of *Rhagoletis indifferens* Curran (Diptera: Tephritidae) in crabapple. Pan-Pac. Entomol. 89, 18–26. 10.3956/2012-48.1

Yee, W.L., Lawrence, T.W., Hood, G.R., Feder, J.L., 2015. New records of *Rhagoletis* Loew, 1862 (Diptera: Tephritidae) and their host plants in western Montana, U.S.A. Pan-Pac. Entomol. 91, 39–57. 10.3956/2014-91.1.039

Yee, W.L., Milnes, J.M., Goughnour, R.B., Bush, M.R., Ray Hood, G., Feder, J.L., 2023. Evidence for adaptation of *Rhagoletis pomonella* (Diptera: Tephritidae) on large-thorn hawthorn, *Crataegus macracantha*, in Okanogan County, Washington State, USA. Environ. Entomol. 52, 455–464. 10.1093/ee/nvad026

