## Supplementary Figures and Tables for "Conserved cold tolerance of *Rhagoletis* species from different host fruits, elevations in Colorado, USA"

```
Rz.ref  TACTTGT--------------------------------------------ATGAAAATTTGCAACTTGTACTAGAAGT
Rp.ref  TACTTGTAAGAAAACTT--ATGATAAGAGGTAATATAAATTTTTTACTTGTATGAAAATTTGCAACTTGTACTAGAAGT
Ha1     TACGAGTTACTGAACTTTTATGATAAGAGGTAATATAAATTTTTTACTTGTATGAAAATTTGCAACTTGTACTAGAAGT
Ha2     TACGAGTTACTGAACTTATTACATAAGAGGTAATATAAATTTTTTACTTGTATGAAAATTTGCAACTTGTACTAGAAGT
Cr1     TACGAGTTACTGAACTTATKACTWGTWTGAAAATATAAATTTTTTACTTGTATGAAAATWWGCAACTTGTACTWGAAGT
Cr2     TACGAGTTACTGAACTTATTACTTGTATGGTAATATAAATTTTTTACTTGTATGAAAATTTGCAACTTGTACTAGAAGT
        ***  ** .  .....            .  ....................********..************.*****
```

**Figure S1**. Alignment of partial internal transcribed spacer (ITS) from two reference (ref) *Rhagoletis zephyria* (Rz) and *Rhagoletis pomonella* (Rp) sequences and four samples from this study (Cr = crabapple, Ha = hawthorn). Rz = ZCO16, Rp = PCO1, obtained from Figure 4 in Smith et al. (2022). Note that the large gap in Rz is not present in our Cr and Ha sequences. Underlined sequence indicates high homology among our samples and Rp reference sequence. Asterisks, same nucleotide at that position in all sequences; dots, same nucleotide in all but one sequence.

**Table S1.** ***Rhagoletis pomonella*** **survival following different durations of exposure to 0°C for between 1 and 16 weeks.** Pupae were chilled for 14weeks at 4°C prior to cold exposure at 0°C. Survival post-cold stress was determined by respirometry and eclosion tracking (completion of development) at 21°C. All pupae were collected from hawthorn fruits in 2018.

| Duration of cold exposure (weeks) | Proportion survived | Proportion eclosed | Mean eclosion time ± s.e. (d) |
|---|---|---|---|
| 1 | 9/9 | 6/9 | 69.7 ± 7.0 |
| 2 | 9/9 | 5/9 | 71.3 ± 4.9 |
| 3 | 9/9 | 5/9 | 53.5 ± 8.9 |
| 4 | 9/9 | 8/9 | 59.5 ± 6.3 |
| 6 | 7/7 | 7/7 | 54.7 ± 3.8 |
| 8 | 9/9 | 8/9 | 58.5 ± 2.5 |
| 10 | 9/9 | 6/9 | 60.4 ± 1.5 |
| 12 | 9/9 | 4/9 | 51.0 ± 4.1 |
| 14 | 7/9 | 5/9 | 47.2 ± 3.7 |
| 16 | 9/9 | 6/9 | 45.5 ± 2.0 |

**Table S2. Kolmogorov-Smirnov (K-S) tests comparing the distribution of post-chill eclosion times in *Rhagoletis pomonella* flies.** *P* values were Bonferroni-corrected for multiple comparisons.

| **Population 1** | **Population 2** | ***D* value** | ***P* value** |
|---|---|---|---|
| Crabapple – Forest Parkway | Crabapple – Congress Park | 0.46038 | < 0.001 |
| Crabapple – Forest Parkway | Hawthorn – Forest Parkway | 0.59365 | < 0.001 |
| Crabapple – Forest Parkway | Hawthorn – East High | 0.66436 | < 0.001 |
| Crabapple – Congress Park | Hawthorn – Forest Rd | 0.23822 | 0.102 |
| Crabapple – Congress Park | Hawthorn – East High | 0.20398 | 0.012 |
| Hawthorn – Forest Parkway | Hawthorn – East High | 0.30835 | 0.002 |
